## Supplementary Material for "Activity-mediated accumulation of potassium induces a switch in firing pattern and neuronal excitability type"

### This PDF file includes:

Figs. S1 to S10.

Tables. S1 to S5.

Eqs. S0.

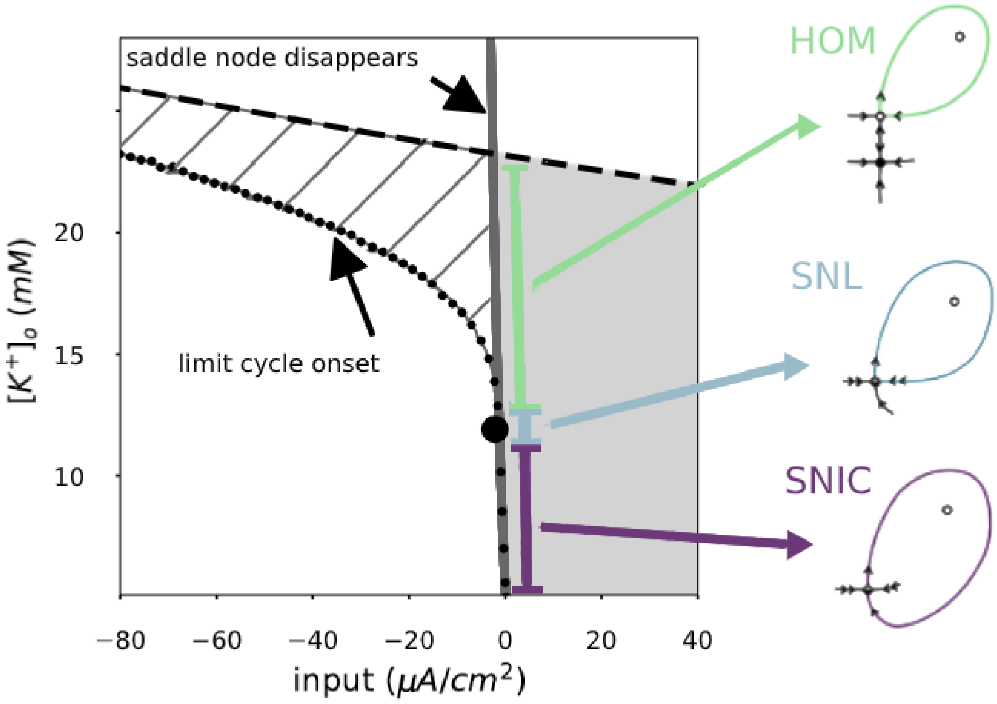


**Fig S1. Transition from rest to spiking (limit cycle onset bifurcations) for different extracellular potassium concentrations.** From bottom to top; SNIC (saddle-node on invariant circle): Purple, SNL (Saddle-node-loop): Blue; HOM (saddle homoclinic orbit): Green. In the SNIC regime the stable node collides with an unstable node, giving rise to a saddle node. The limit cycle orbit passes through the saddle node, the trajectory leaves the saddle node along the semi-stable manifold. After one period trajectory approaches the saddle node along the same semi-stable manifold. At the SNL point, trajectories leave the saddle node along the semi-stable manifold as in the SNIC case, but after one period those trajectories approach the saddle node along the strongly stable manifold. Notice that the SNL orbit is smaller than the SNIC orbit, and has a shorter period. In the HOM regime a stable node and a limit cycle coexist. External perturbations shift the state of the system from the stable node to the attraction domain of the limit cycle attractor.

$$I_{\text{pump}} = \begin{cases} 0 & [\text{Na}^+]_i \leq [\text{Na}]_s \\ \frac{I_{\text{maxp}}}{1+\exp(k_{\text{Na}}([\text{Na}^+]_i - [\text{Na}]_s))} \frac{2}{1+\exp(-K_s([\text{K}^+]_o - [\text{K}^+]_s))} & [\text{Na}^+]_i > [\text{Na}]_s \end{cases} \tag{S0}$$

| | $\tau_{fast}$ $[ms]$ | $\tau_{slow}$ $[ms]$ | D$_{fast}$ $[mV]$ | D$_{slow}$ $[mV]$ | D$_{ss}$ $[mV]$ |
|---|---|---|---|---|---|
| count | 48.000000 | 48.000000 | 48.000000 | 48.000000 | 4.800000e+01 |
| mean | 480.314157 | 17728.034470 | 5.906181 | 20.379248 | 2.352945e+01 |
| std | 676.504019 | 7919.924120 | 2.470383 | 6.366880 | 1.147889e+01 |
| min | 61.687083 | 4312.710113 | 0.000001 | 7.860144 | 6.323943e-07 |
| 25% | 170.353968 | 12050.396908 | 4.334064 | 17.144954 | 1.672852e+01 |
| 50% | 232.372743 | 16797.597216 | 5.823583 | 20.102315 | 2.359666e+01 |
| 75% | 368.297499 | 21156.877421 | 7.097274 | 22.357364 | 3.093022e+01 |
| max | 3083.939293 | 45815.441228 | 14.308320 | 48.164289 | 5.970884e+01 |

**Table S1. Summary of the distribution of the best fit of the parameters for each of the 48 traces. Depolarizing pulses applied at a 40Hz rate**. See

| | $\tau_{fast}$ $[ms]$ | $\tau_{slow}$ $[ms]$ | D$_{fast}$ $[mV]$ | D$_{slow}$ $[mV]$ | D$_{ss}$ $[mV]$ |
|---|---|---|---|---|---|
| count | 73.000000 | 73.000000 | 73.000000 | 73.000000 | 73.000000 |
| mean | 935.652651 | 15707.653647 | 5.064957 | 7.737310 | 40.589393 |
| std | 599.644611 | 9936.369466 | 2.492782 | 4.081336 | 7.589756 |
| min | 125.198826 | 1485.350564 | 0.965347 | 0.000004 | 0.000024 |
| 25% | 466.154928 | 10033.043848 | 3.045424 | 5.644995 | 36.885021 |
| 50% | 745.856489 | 13876.630769 | 4.882892 | 7.078889 | 42.152299 |
| 75% | 1319.313755 | 18294.743647 | 6.750978 | 8.840077 | 45.033475 |
| max | 2436.623039 | 52548.189654 | 11.642939 | 32.905853 | 52.764149 |

**Table S2. Summary of the distribution of the best fit of the parameters for each of the 73 traces. Hyperpolarizing pulses.**

| Gating dynamics | |
|---|---|
| $\frac{dm_{Na}}{dt}$ | $\alpha_m (1 - m_{Na}) - \beta_m m_{Na}$ |
| $\frac{dh_{Na}}{dt}$ | $\alpha_h (1 - h_{Na}) - \beta_h h_{Na}$ |
| $\frac{dn_K}{dt}$ | $\alpha_n (1 - n_K) - \beta_n n_K$ |

**Table S3. Gating dynamics used for the excitable portion of the model.**

| Functions | |
|---|---|
| $E_K$ | $\frac{RT}{F} \log\left(\frac{K_o}{K_i}\right)$ |
| $E_L$ | $\frac{RT}{F} \log\left(\frac{K_o P_K + Na_o P_{Na}}{K_i P_K + Na_i P_{Na}}\right)$ |
| $E_{Na}$ | $\frac{RT}{F} \log\left(\frac{Na_o}{Na_i}\right)$ |
| $\alpha_h$ | $\frac{0.128 q_h e^{-\frac{\alpha_{hV}}{18} - \frac{v}{18mV}}}{ms}$ |
| $\beta_h$ | $\frac{4 q_h}{ms\left(e^{-\frac{\beta_{hV}}{5} - \frac{v}{5mV}} + 1\right)}$ |
| $\alpha_m$ | $\frac{q_m\left(0.32\alpha_{mV} + \frac{0.32v}{mV}\right)}{ms\left(1 - e^{-\frac{\alpha_{mV}}{4} - \frac{v}{4mV}}\right)}$ |
| $\beta_m$ | $\frac{q_m\left(0.28\beta_{mV} + \frac{0.28v}{mV}\right)}{ms\left(e^{\frac{\beta_{mV}}{5} + \frac{v}{5mV}} - 1\right)}$ |
| $\beta_n$ | $\frac{0.5 q_n e^{-\frac{\beta_{nV}}{40} - \frac{v}{40mV}}}{ms}$ |

**Table S4. Expressions used for the excitable portion of the model.**

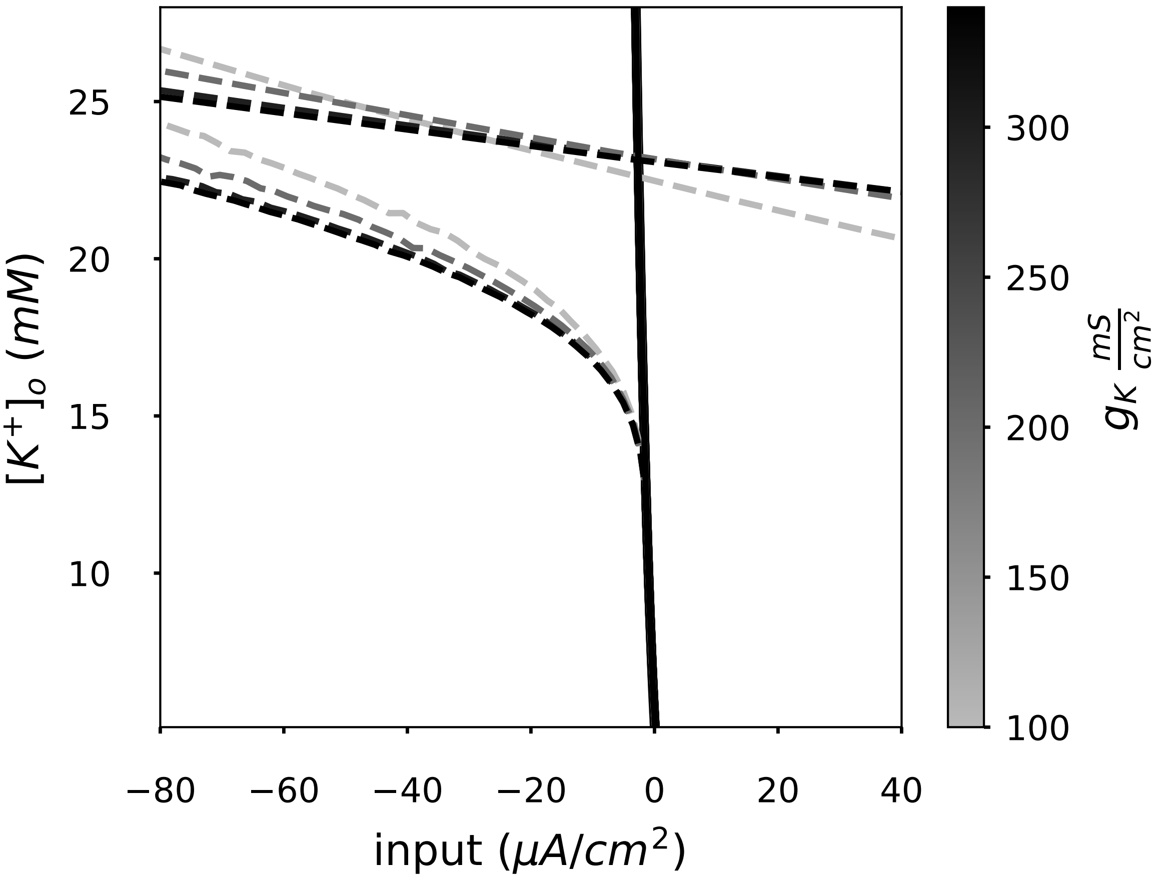


**Fig S2. Changes in the conductance of the delayed rectifier potassium current ($g_K$) distorts the bistable region portrayed in Fig.3A.** Same bifurcation diagram portrayed in Fig.3A for different $g_K$. Here the curves correspond to the delayed rectifier conductance of $g_K$; 100, 200, 300, and 340 $msiemens/cm^2$. As $g_K$ increases, the limit cycle onset, and the depolarization block lines are shifted towards higher extracellular potassium concentrations.

| Parameters excitable portion | |
|---|---|
| $g_K$ | $\frac{200mS}{cm^2}$ |
| $g_L$ | $\frac{0.1mS}{cm^2}$ |
| $g_{Na}$ | $\frac{100mS}{cm^2}$ |
| $C$ | $\frac{1.0uF}{cm^2}$ |
| $\alpha_{hV}$ | 50 |
| $\alpha_{mV}$ | 54 |
| $\alpha_{nV}$ | 52 |
| $\beta_{hV}$ | 27 |
| $\beta_{mV}$ | 27 |
| $\beta_{nV}$ | 57 |

**Table S5. Parameters used for the excitable portion of the model.**

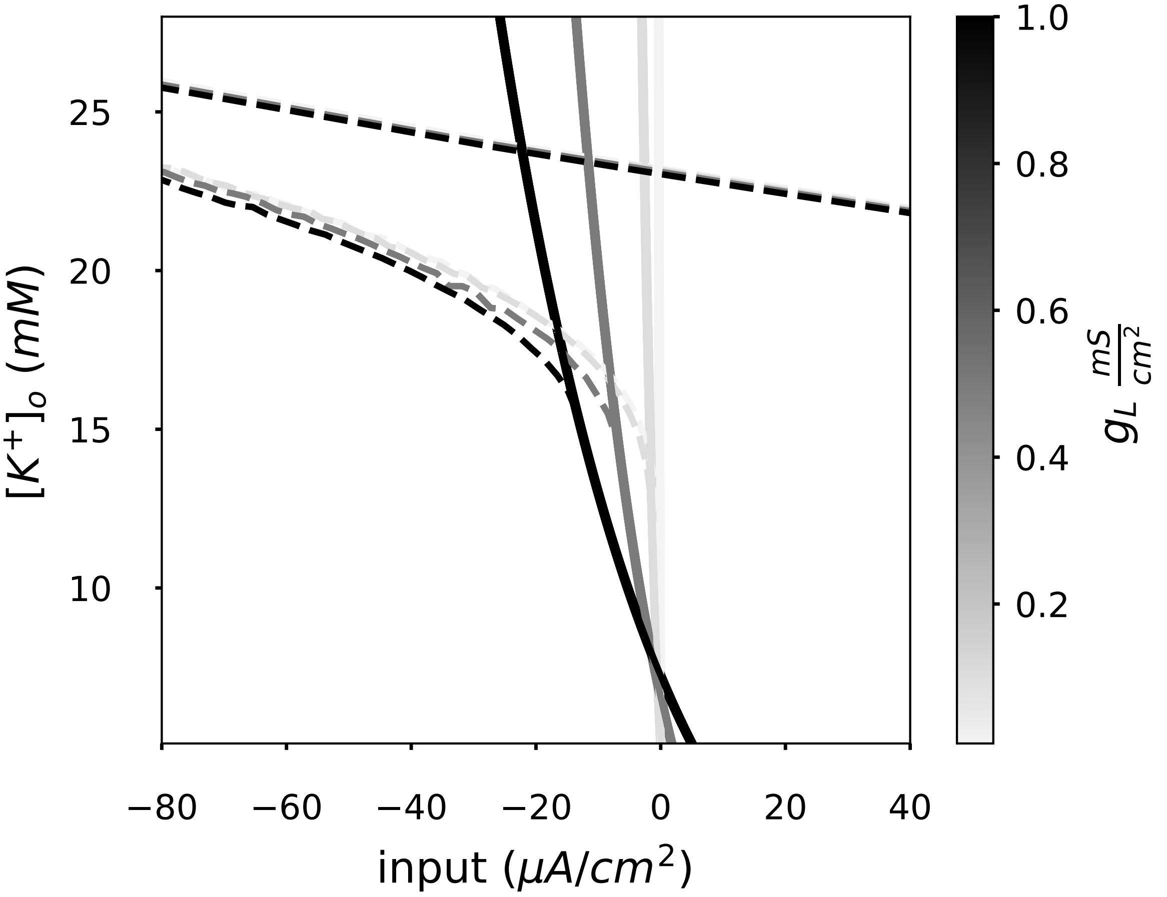


**Fig S3. Changes in the leak conductance ($g_L$) distorts the bistable region portrayed in Fig.3A.** Same bifurcation diagram portrayed in Fig.3A for different $g_L$. Here the curves correspond to leak conductances of $g_L$; 0.01, 0.1, 0.5, and 1.0 $msiemens/cm^2$. As $g_L$ increases, the bistable region is shifted towards higher extracellular potassium concentrations. Another effect of more leaky neurons, is that the dependence of the spiking threshold on extracellular potassium is more prominent.

| Parameters ionic concentration dynamics | |
|---|---|
| $\rho$ | $\frac{4000}{cm}$ |
| $F$ | $\frac{96484.6C}{mol}$ |
| $\frac{Vol_i}{Vol_e}$ | 0.2 |
| $I_{max}$ | $\frac{40.0uA}{cm_2}$ |
| $[Na^+]_o$ | $140mM$ |
| $Na_s$ | $\frac{0.1}{mM}$ |
| $K_{Na}$ | $20mM$ |

**Table S6. Parameters used for the ionic concentration dynamics portion of the model.**

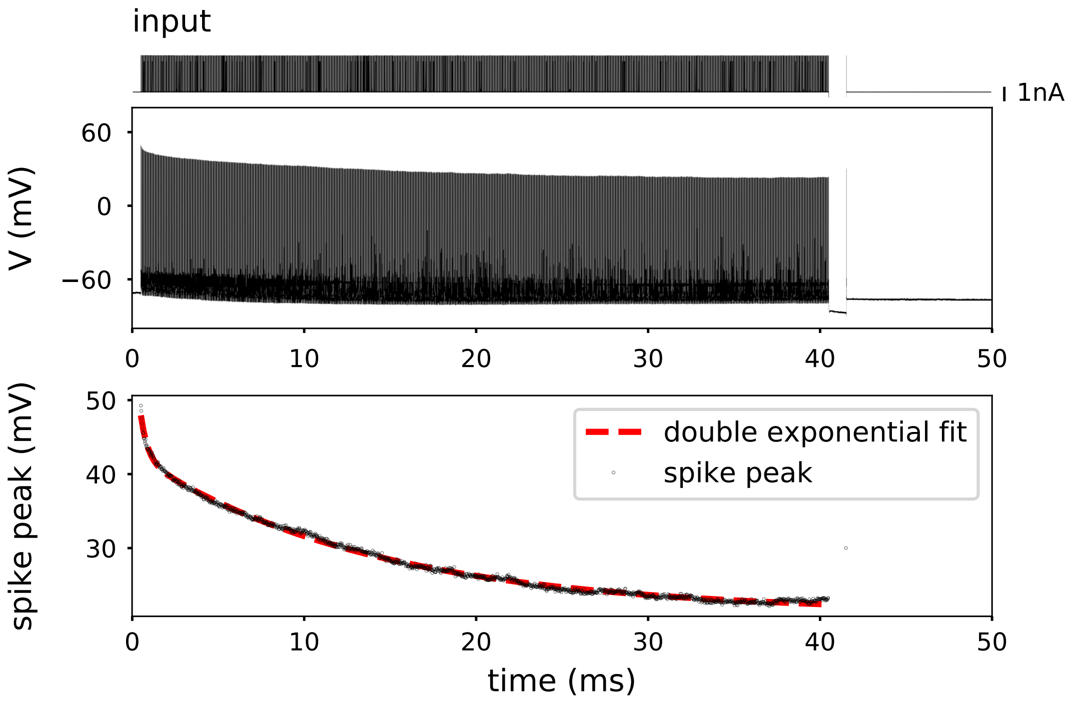


**Fig S4. Slow decay of spike amplitudes. Voltage recording of a neuron experiencing depolarizing pulses applied at 40 Hz. The fast and slow time constants of amplitude decay were $\tau_{fast} = 410(ms)$ and $\tau_{slow} = 13.6(sec)$, respectively..** Notice that the peak of the last spike fails to recover to the initial amplitude after the one-second-long hyper-polarizing pulse.

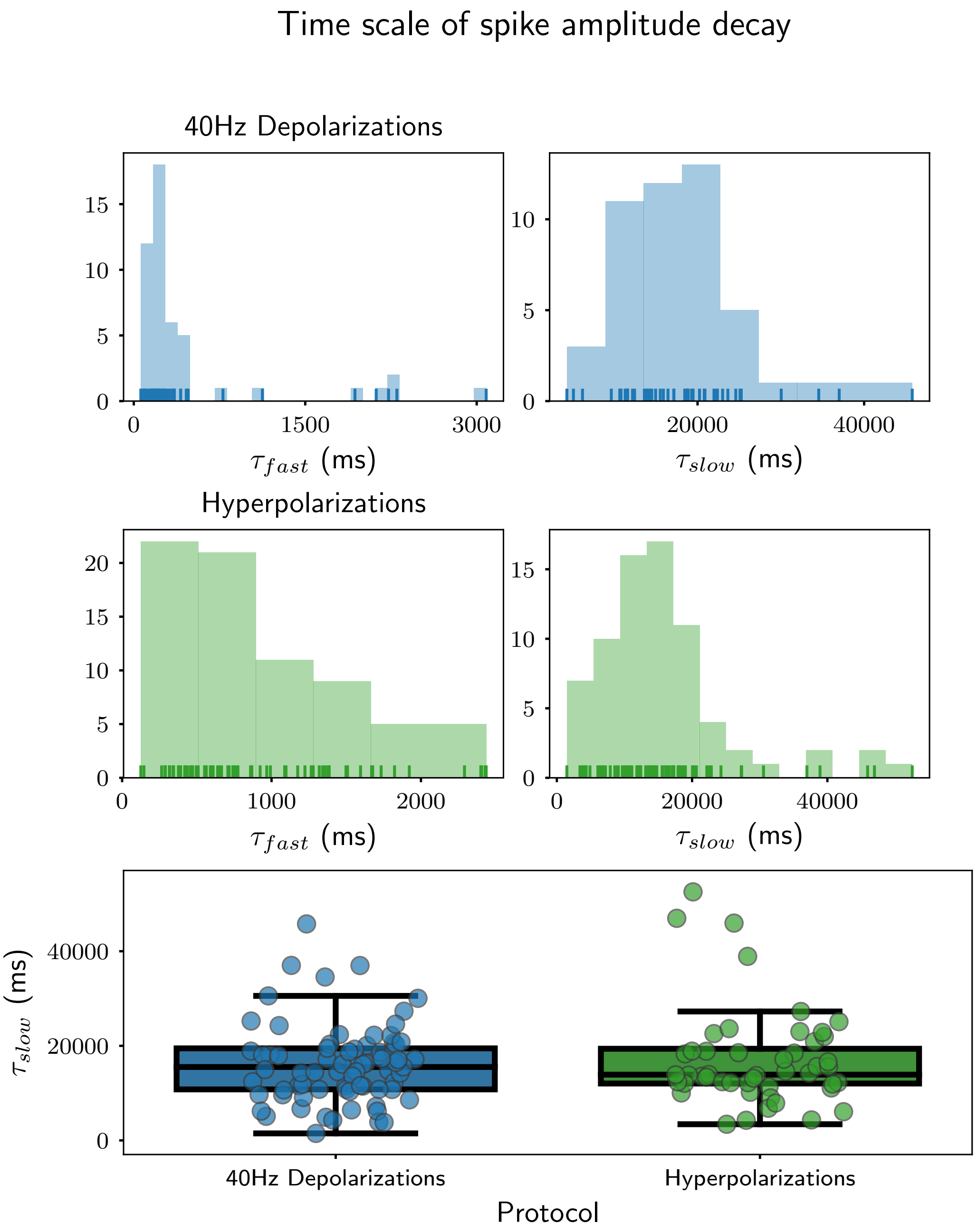


**Fig S5. Distribution of time scales of the double exponential decay (equation ??) of the spike amplitude.** Two protocols were used to measure the time scales of spike amplitude decay, an example of the "40 Hz Depolarizations" is shown in Fig. S4, and an example of the "Hyperpolarization" is shown in Fig. 5 Notice that the distribution of $\tau_{slow}$ is independent of the protocol used.

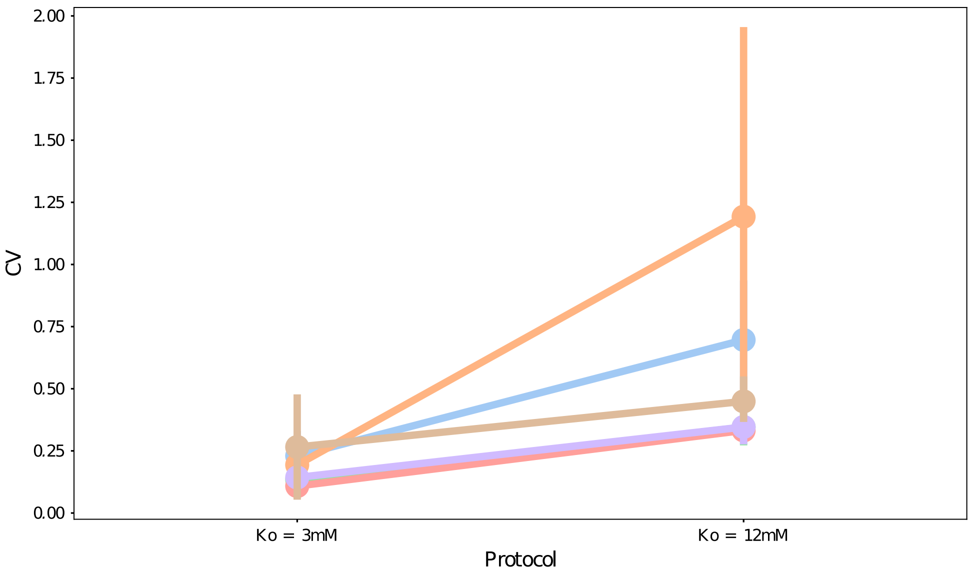


**Fig S6. Spiking variability calculated as the coefficient of variation ($CV = \frac{\sigma}{\mu}$) for all cells sampled, when stimulating with white noise added to the baseline input (n=6).** An increase from 3 mM to 12 mM in extracellular potassium increased the spiking variability of 5 out of 6 cells measured.

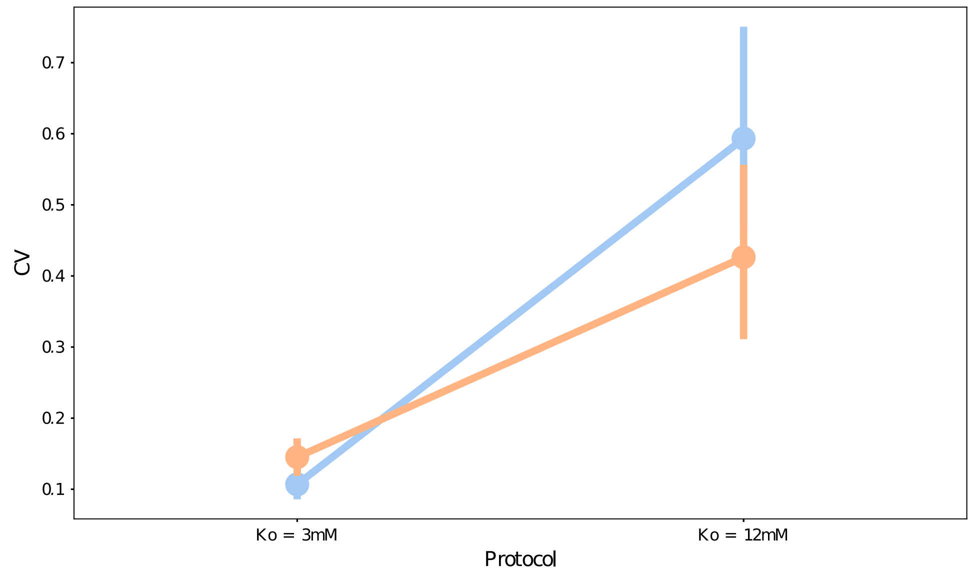


**Fig S7. Spiking variability calculated as the coefficient of variation ($CV = \frac{\sigma}{\mu}$) for all cells sampled, when stimulating with baseline input (n=2).** An increase from 3 mM to 12 mM in extracellular potassium increased the spiking variability of 2 out of 2 cells measured. The main source of stimuli irregularity was the network activity.

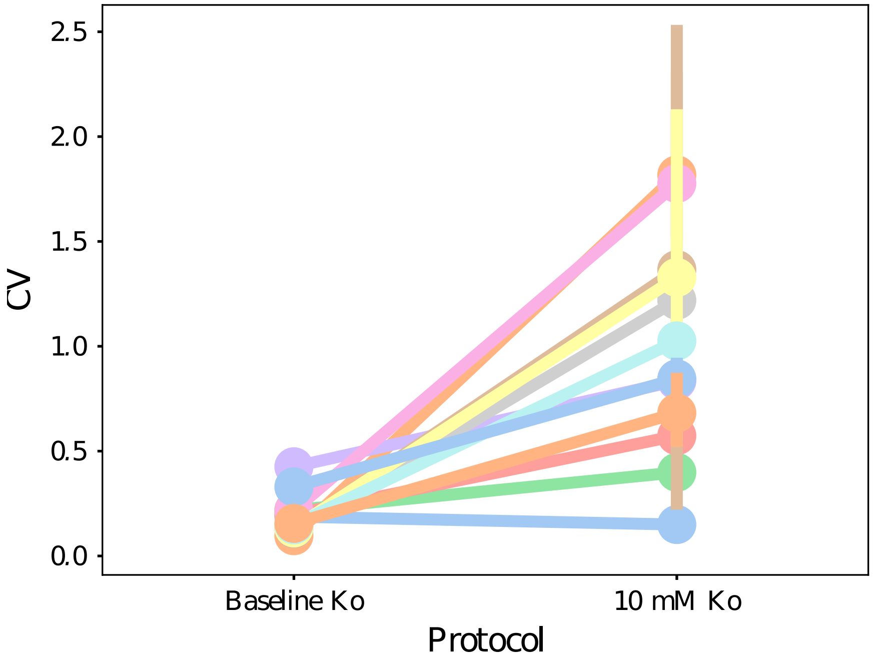


**Fig S8. Spiking variability calculated as the coefficient of variation ($CV = \frac{\sigma}{\mu}$) for all cells sampled after blocking synaptic input, under baseline input stimulation (n=12).** An increase from 3 mM to 10 mM in extracellular potassium increased the spiking variability of 8 out of 12 cells measured.

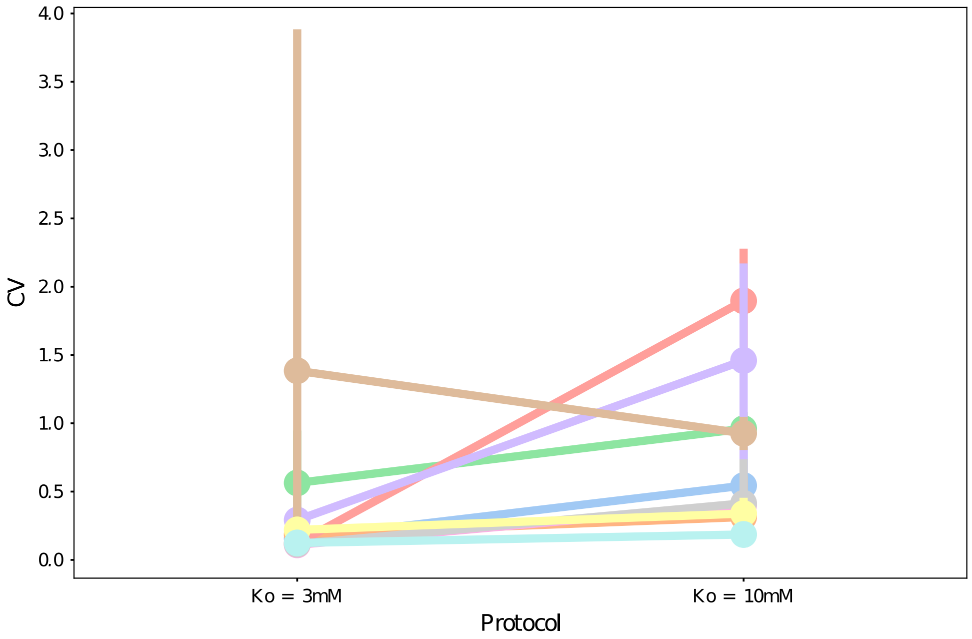


**Fig S9. Spiking variability calculated as the coefficient of variation ($CV = \frac{\sigma}{\mu}$) for all cells sampled (n=10).** An increase from 3 mM to 10 mM in extracellular potassium increased the spiking variability of 3 out of 10 cells measured. The main source of stimuli irregularity was the network activity.

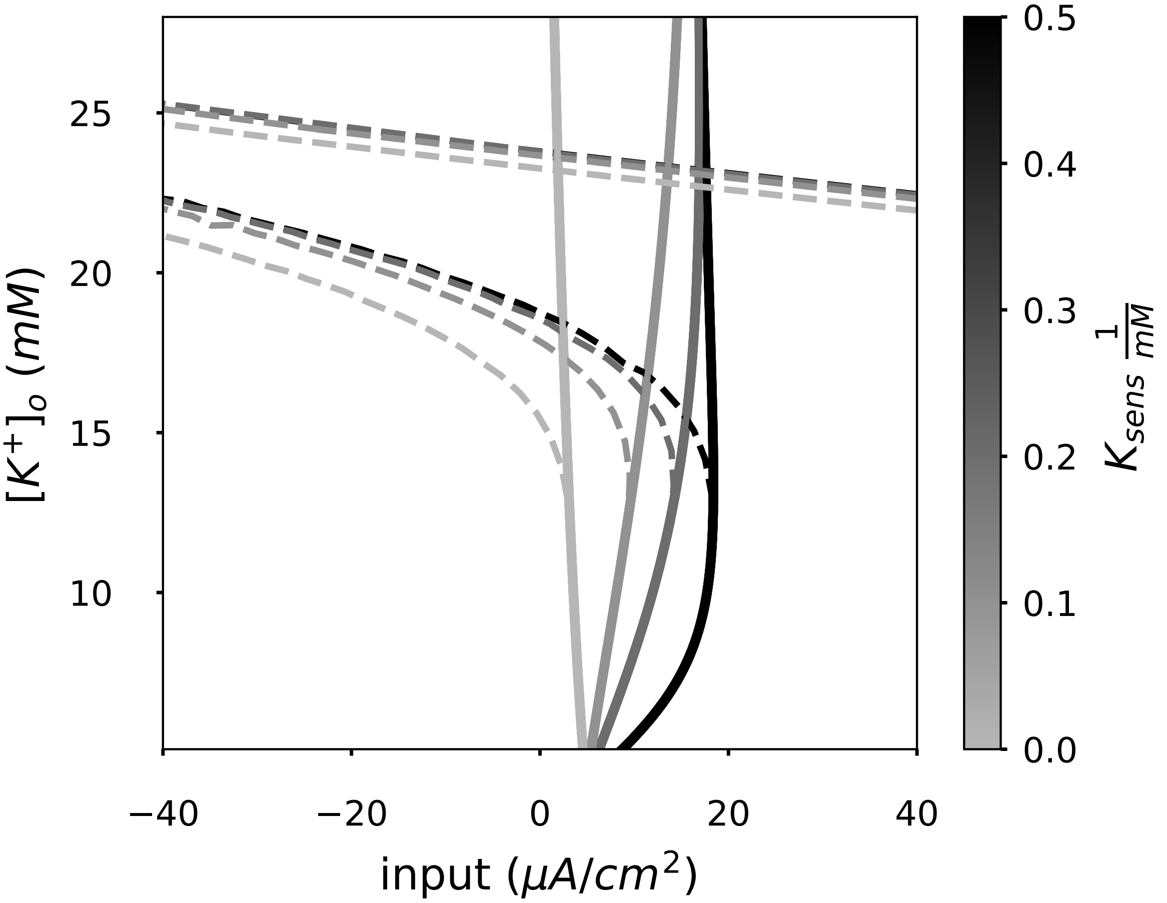


**Fig S10. Extracellular potassium and $[K^+]_o$ pump's sensitivity ($K_{\mathrm{sens}}$) dependent bistable area.** Same bifurcation diagram portrayed in Fig. 6 for different $[K^+]_o$ pump's sensitivity. Here 0,0.1,0.2 and 0.5 $1/mM$ sensitivities to $[K^+]_o$ ($K_{\mathrm{s}}$) are portrayed and $[\mathrm{K}^+]_s$ is fixed to 4 mM for all curves, the expression of the pump that was used here resembles isoform $\alpha_2$ (eq. S0 ). $K_{\mathrm{s}}$ distorts the saddle node bifurcation line, curving it towards more depolarized currents, i.e shifting the spiking threshold towards higher input currents.
